# Quantifying E2F1 protein dynamics in single cells

**DOI:** 10.1101/634956

**Authors:** Bernard Mathey-Prevot, Bao-Tran Parker, Carolyn Im, Cierra Hong, Peng Dong, Guang Yao, Lingchong You

## Abstract

The Rb/E2F pathway plays a central role in regulating cell-fate decisions and cell-cycle progression. The E2F1 protein, a major effector of the pathway, is regulated via a combination of transcriptional, translational and posttranslational constraints. Elucidating the regulation and impact of the Rb/E2F pathway requires direct measurement of E2F1 dynamics in single cells. To this end, we have engineered fluorescent E2F1 protein reporters to enable live detection and quantification in single cells. The reporter constructs expressed an E2F1-Venus fusion protein under the regulation of the mouse or human E2F1 promoter and contained or excluded the 3’UTR of the E2F1 gene, a sequence that contains miRNA regulatory regions that modulate expression of the protein. Expression of the reporter protein was highly dynamic during the cell cycle: there was no or little fluorescent signal in G_0_, but levels steadily increased during late G_1_ and peaked during mid to late S phase before returning to baseline before the onset of mitosis. The absence of the E2F1 3’UTR in the constructs led to considerably higher steady-state levels of the fusion protein, which although normally regulated, exhibited a slightly less complex dynamic profile during the cell cycle or genotoxic stress. Lastly, the presence or absence of Rb failed to impact in substantial ways the overall detection and levels of the reporters.

## Introduction

E2Fs are a family of transcription factors that orchestrate the traverse of the G_1_/S and G_2_/M phases of the cell cycle by regulating the expression of critical genes promoting proper cell cycle progression (Fischer and Müller 2017; Bertoli, Skotheim, and de Bruin 2013; Ren et al. 2002). This family of transcription factors has been subdivided into both activators (E2F1-E2F3a) and repressors (E2F3b, E2F4-E2F8) (van den Heuvel and Dyson 2008). Rb, a member of the pocket protein family negatively regulates the function of E2F activators by directly binding to them in a cell cycle-dependent manner (Henley and Dick 2012). Its inhibition is specifically relieved during the G1 phase through hyper-phosphorylation by G_1_ cyclins/CDK complexes (Sherr and Roberts 2004; Henley and Dick 2012) Loss of Rb or deregulation of targets of E2F transcription factors have been associated with numerous human malignancies, highlighting the importance of this pathway in cell cycle regulation and disease (Chen, Tsai, and Leone 2009).

To better understand how the dynamics of activation of the Rb/E2F pathway correlate with quiescence and proliferation of cells exposed to growth stimuli, we had developed two reporters that captured the transcriptional activation of E2F1 and E2F activity respectively (Dong et al. 2014, 2018). However, E2Fs are further regulated at the post-translational level, resulting in dynamically regulated amounts of those proteins during the cell cycle. Although this aspect of E2F1 expression was reflected by the behavior of the E2F activity reporter, it fell short of providing a comprehensive and direct picture of the dynamic changes in E2F1 protein in a single cell. As the overall balance of E2F repressors to activators can affect cellular outcomes (e.g., proliferation, differentiation, senescence, or apoptosis (Dimova and Dyson 2005; Dimri, Itahana, and Acosta 2000)), there was a need to complement our existing reporters with a third reporter construct which would emulate the specific behavior of the E2F1 protein under various conditions.

## Results

### Design rationale of the E2F1 protein reporter constructs

Building on our experience with our two previous reporters (Fig 1A), we defined a set of criteria for the E2F1 protein reporter: 1) it should be subject to the same transcriptional regulation as the endogenous E2F1 gene, as was the case for our earlier reporters (Dong et al. 2014, 2018); 2) the influence of the E2F1 3’UTR on the expression of the reporter protein should be evaluated, as this region is targeted by miRNAs shown to regulate the levels of E2F1 protein (O’Donnell et al. 2005; Pickering, Stadler, and Kowalik 2009); 3) ectopic expression of the reporter protein should not perturb the overall E2F activity in a cell to avoid altering the balance between EF2 activators and repressors that dictate different cell fate decisions (Iaquinta and Lees 2007; Poppy Roworth, Ghari, and La Thangue 2015) and 4)tThe reporter protein should be fluorescent for purpose of live detection and contain all E2F1 residues known to be subject of post-translational modification, to maximize our ability to capture the dynamic expression of E2F1 under different experimental conditions (Poppy Roworth, Ghari, and La Thangue 2015).

**Figure 1:**
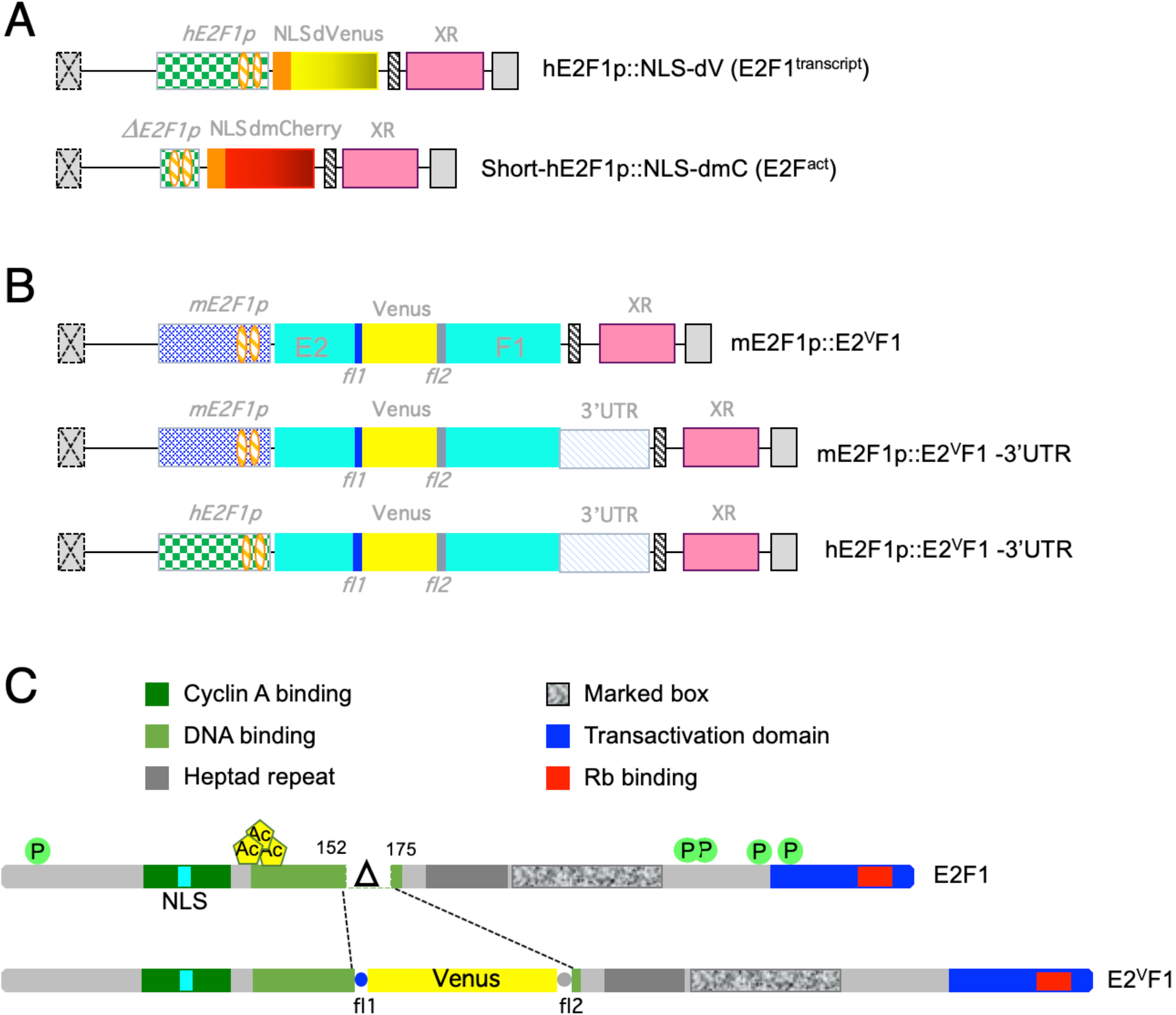
E2F1 protein reporter constructs. **(A)** Schematics of the E2F1 transcriptional and E2F activity reporters previously published (Dong et al. 2014, 2018). *hE2F1p*: human E2F1 promoter; Δ *E2F1p*: short human E2F1 promoter; NLS: SV40 nuclear localization domain; dVenus: destabilized Venus; dmCherry: destabilized monomeric Cherry; XR: drug resistance gene (puromycin or neomycin); orange hatched ovals: E2F consensus binding sites. **(B)** Schematic of E2F1 protein reporters generated in the pQCXIP (or pQCXIN) expression construct (Takara Bio/Clontech). *mE2F1p* and *hE2F1p*: E2F1 mouse and human promoter, respectively (Dong et al. 2014); E2: sequence encoding a.a. 1-152 of hE2F1; *Venus*: sequence coding for the Venus fluorescent protein; F1: sequence encoding a.a. 175-437 of hE2F1; 3’UTR: 3’ untranslated region of hE2F1 gene. **(C)** Schematics of human E2F1 and E2^V^F1 proteins. Main functional domains are highlighted. P: phosphorylation sites; Ac: acetylation sites; NLS: nuclear localization domain; Δ : amino acids 153-174 deletion corresponding to leucine zipper in DNA binding domain (LNWAAEVLKVQKRRIYDITNVL); fl1: flexible linker 1; fl2: flexible linker 2.

To satisfy these criteria, we decided to use the previously validated mouse or human E2F1 promoter (Dong et al. 2014, 2018) and include or omit the E2F1 3’UTR region in our constructs (Fig. 1B). In addition, the E2F1 protein reporter constructs were designed to encode a fusion protein (E2^V^F1) of 686 amino acids (a.a.) (Fig. 1C), consisting of the N-terminal region of human E2F1 (a.a. 1-152) fused to the fluorescent protein Venus flanked at either end with a flexible peptide linker, and followed by the rest of the E2F1 C-terminal region (a.a. 175-437). In the process, a small region of E2F1 corresponding to the winged-helix DNA binding domain was deleted (a.a. 152-174), with the intent of preventing the reporter protein to be transcriptionally active (Zheng et al. 1999)). Other than the deleted residues, which have not been described to be targeted by post-translational modifications, E2^V^F1 retains all E2F1 residues reported targeted for post-translational regulation during the cell cycle or in response to stress conditions (Fig. 1C) (Munro, Carr, and La Thangue 2012).

### Expression of the E2^V^F1 fusion protein

Viral stocks corresponding to our constructs were used to infect rat and human fibroblasts (Fig. 2A and 2B, respectively) or human mammary epithelial cells (Fig. 2C). After selection in puromycin, polyclonal populations were either used directly or subjected to limiting dilution to isolate single cell clones. In all cases, expression of the reporter protein was detected in the form of two bands, which were absent in uninfected cells (Fig. 2). The top band corresponds to the expected size of the fusion protein (686 a.a., ∼73 kDa). As the observed difference in migration (∼19 kDa) is too large to be caused by post-translational modifications, the lower, the lower band likely represents a truncation product of the mature form. However, we cannot rule out that it corresponds to the translation product of E2^V^F1 at an internal start site, perhaps at the initiating methionine codon of Venus which, together with the surrounding nucleotides, conforms to a strong Kozak consensus sequence (Kozak 1987). The steady-state levels of E2^V^F1 expression appear to be higher than that of endogenous E2F1 in all cell types tested (SupFig. 1A). Furthermore, there was a 4-8 fold increase in the steady-state amount of E2^V^F1 protein in REF52^E2vF1^ clone 8 cells compared to the levels observed in REF52 clones transduced with the same construct but containing the E2F1 3’UTR (Fig. 2A, lane d vs. lane b or c). This increase was not unique to clone 8 cells but was also observed in the REF52 parental polyclonal population of cells transduced with the E2^V^F1 construct and other single cell clones isolated from that population (data not shown). In contrast, all cells transduced with versions of the E2^V^F1-3’UTR construct expressed the fusion protein at similar or lower levels to that detected in REF52^E2vF1-3’UTR^ clone B or D cells (Figure 2B and SupFig.1). As such, the presence or absence of the 3’UTR in our protein reporter constructs appears to influence the amount of the fusion protein expressed in cells (Fig. 2).

**Figure 2:**
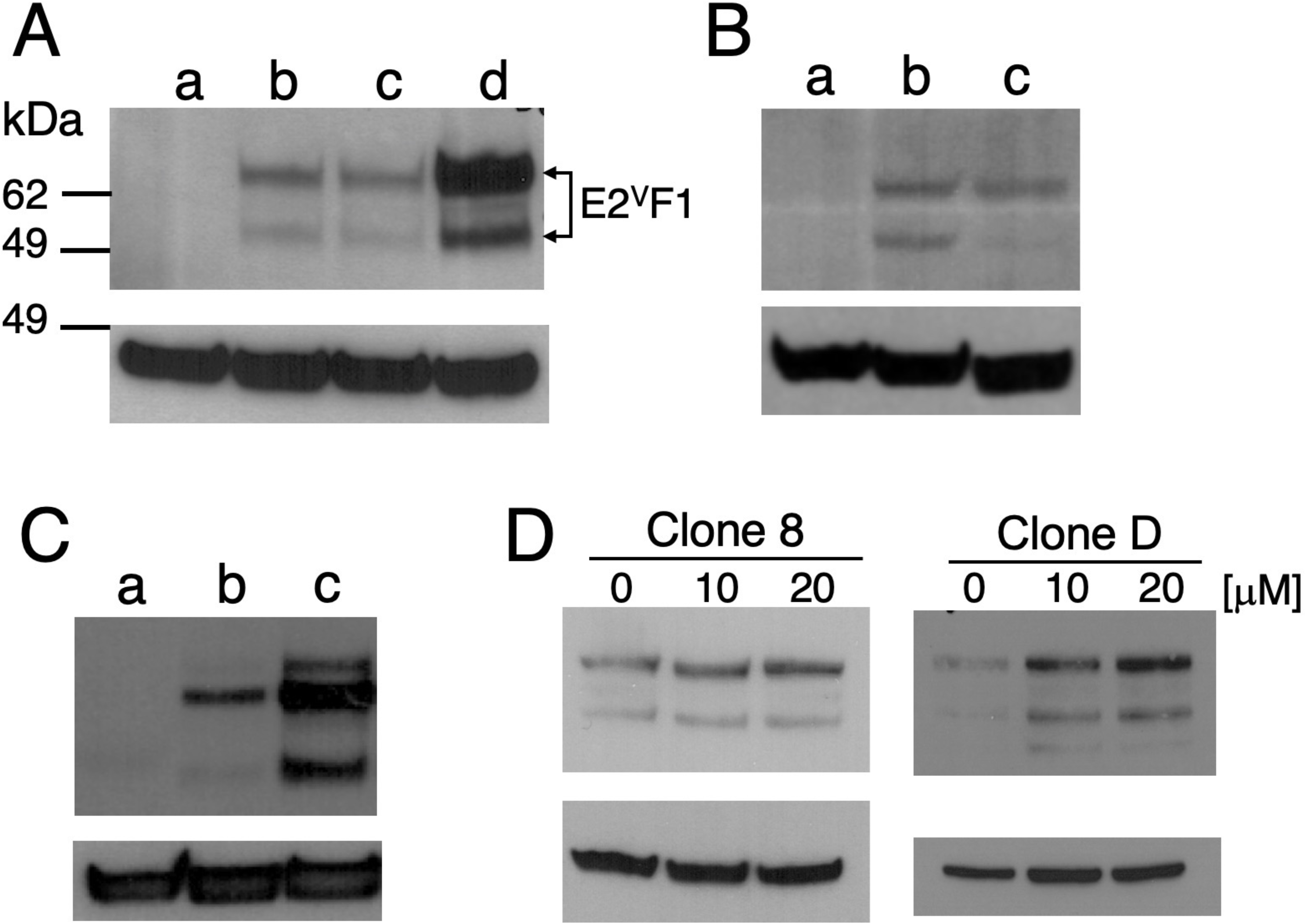
Characterization of the E2^V^F1 fusion protein. (**A - D)** Cell extracts were prepared from cells grown under normal or experimental conditions (as indicated) and used for detection of E2^V^F1 protein: Top panel: α -E2F1 antibody. Bottom panel: β -actin antibody. **(A)** a: REF52 (uninfected); b: REF52^E2vF1-3’UTR^ (clone B); c: REF52^E2vF1-3’UTR^ (clone D); d: REF52^E2vF1^ (clone 8). **(B)** a: Wi-38 (uninfected); b: WI-38^E2vF1-3’UTR^ (polyclonal); c: REF52^E2vF1-3’UTR^ (clone D). (**C)** a: HME^E2Fact^ (clone 1) cells grown in full medium. b: HME^E2Fact^ Rb+, E2^V^F1-3’UTR (clone R15) cells were incubated sequentially in starvation medium #1 and #2 for a total of 48h (see Material and Methods). c: HME^E2Fact^ Rb+, E2^V^F1-3’UTR (clone R15), starved for 48h and grown again with full medium for 24h. (Top panel: α -E2F1 antibody. Bottom panel: β -actin antibody). **(D)**. Extracts from REF52^E2vF1^ (clone 8) or REF52^E2vF1-3’UTR^ (clone D) grown in the absence or in the presence of increasing amount of cisplatin [μ M].

To further validate our E2F1 protein reporter, we tested whether expression of E2^V^F1 protein responded in a similar fashion to various experimental conditions known to affect the steady-state level of the endogenous E2F1 protein. It is well established that expression levels of E2F1 protein and mRNA dramatically decrease in response to serum starvation (Wong et al. 2011). Cultures of HME^E2Fact^ Rb+, E2^V^F1-3’UTR (clone R15) cells were grown in minimal medium lacking growth supplement additives (see Material and Methods). Cells were either lysed after 48h or released back into the cell cycle by adding fresh full medium for 24h. Cell extracts from both conditions were analyzed for the expression of E2^V^F1. We observed very little expression of E2^V^F1 protein in cells arrested in G_0_, whereas cultures released into the cell cycle for about 24h expressed similar amounts of the fusion protein as compared to cultures grown continuously in full medium (Fig. 2C). As genotoxic stress results in E2F1 protein stabilization (Lin, Lin, and Nevins 2001), we tested the effect of cisplatin on E2^V^F1 protein expression. Cell extracts from REF52^E2vF1^ (clone 8) and REF52^E2vF1-3’UTR^ (clone D) cells treated with cisplatin contained increased amount of E2^V^F1 in a dose-dependent manner compared to control treated cells (Fig. 2D). The effect was more prominent in clone D cells, as these cells express a lower basal level of E2^V^F1(Fig. 2A, lane c) under normal conditions.

Having established that the fusion protein E2^V^F1 recapitulated the behavior of endogenous E2F1 protein under different experimental conditions, we next characterized its localization. All cells transduced with the reporter protein constructs exhibited detectable fluorescent signals (Fig. 3). A stronger signal was observed in cells transduced with the construct lacking the 3’UTR (Sup Fig. 2), consistent with the higher steady-state amounts of E2^V^F1 associated with this construct (Figure 2A). As is the case for E2F1 (Magae et al. 1996; Allen et al. 1997; Ivanova, Vespa, and Dagnino 2007), localization of the fluorescent signal corresponding to E2^V^F1 is largely restricted to the nucleus (Fig. 3). The cell-to-cell variability in the E2^V^F1 signal in unsynchronized clone 8 cells reflects the dynamic modulation of protein expression during the cell cycle (Fig. 3A).

**Figure 3:**
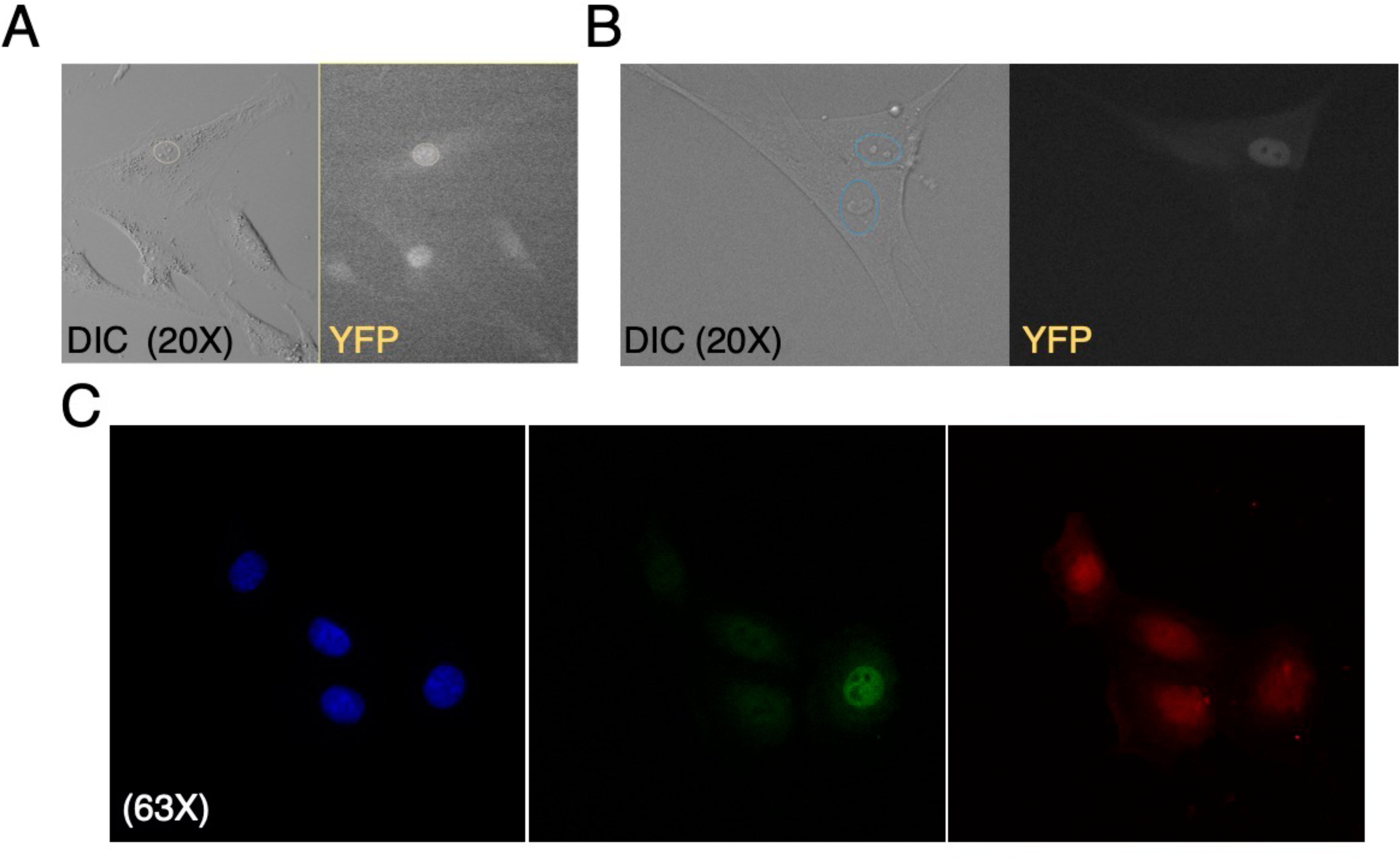
Live detection and nuclear localization of E2^V^F1. Rat or human fibroblasts were grown in 35 mm Mattek optic plates (A and B) for live imaging, and HME cells were grown on glass coverslips (C) placed in regular 35 mm tissue culture dishes before being fixed and permeabilized for immunofluorescence detection. **(A)** Live REF52^E2vF1^ (clone 8) cells imaged under DIC and YFP illumination respectively (Olympus VivaView FL microscope, 20X). Yellow, dotted-line oval highlights nucleus in top cell in DIC panel. **(B)** Live WI-38^E2vF1-3’UTR^ (polyclonal) cells imaged as in A. Cyan, dotted ovals highlight nuclei in the two cells shown in DIC panel. **(C)** Confocal images of fixed HME^E2Fact^ Rb-, E2^V^F1-3’UTR (polyclonal) cells, taken in the DAPI, EYFP and mcherry channels respectively (Zeiss 780 inverted confocal microscope, 63X/1.4 NA Oil Plan-Apochromat DIC).

### Dynamic expression of E2^V^F1 during the cell cycle

To quantify the temporal dynamics of E2^V^F1 during the cell cycle, we initially focused on REF52^E2vF1^ clone 8 cells as they expressed the strongest signal (Fig. 3, SupFig. 2). Clone 8 cells were serum starved for 48h and then released back into the cell cycle by addition of full medium containing 10% serum. Live images were taken every 30 min over 36h and the signal intensity was quantified for each time point up to the time of the first cell division, and occasionally for longer in one the daughter cells. Although E2^V^F1 fluorescence intensity was variable among different single cells at any given time (SupFig. 3), it remained low in each cell over a few hours after serum addition before sharply increasing to a maximum and then decreasing to low or background levels shortly before or at the time of mitosis (Fig. 4). The dynamic behavior of the reporter protein observed in single cells agree with population measurements of endogenous E2F1 protein we had previously observed in REF52 cell extracts (see SupFig. 1b in (Dong et al. 2014)).

**Figure 4:**
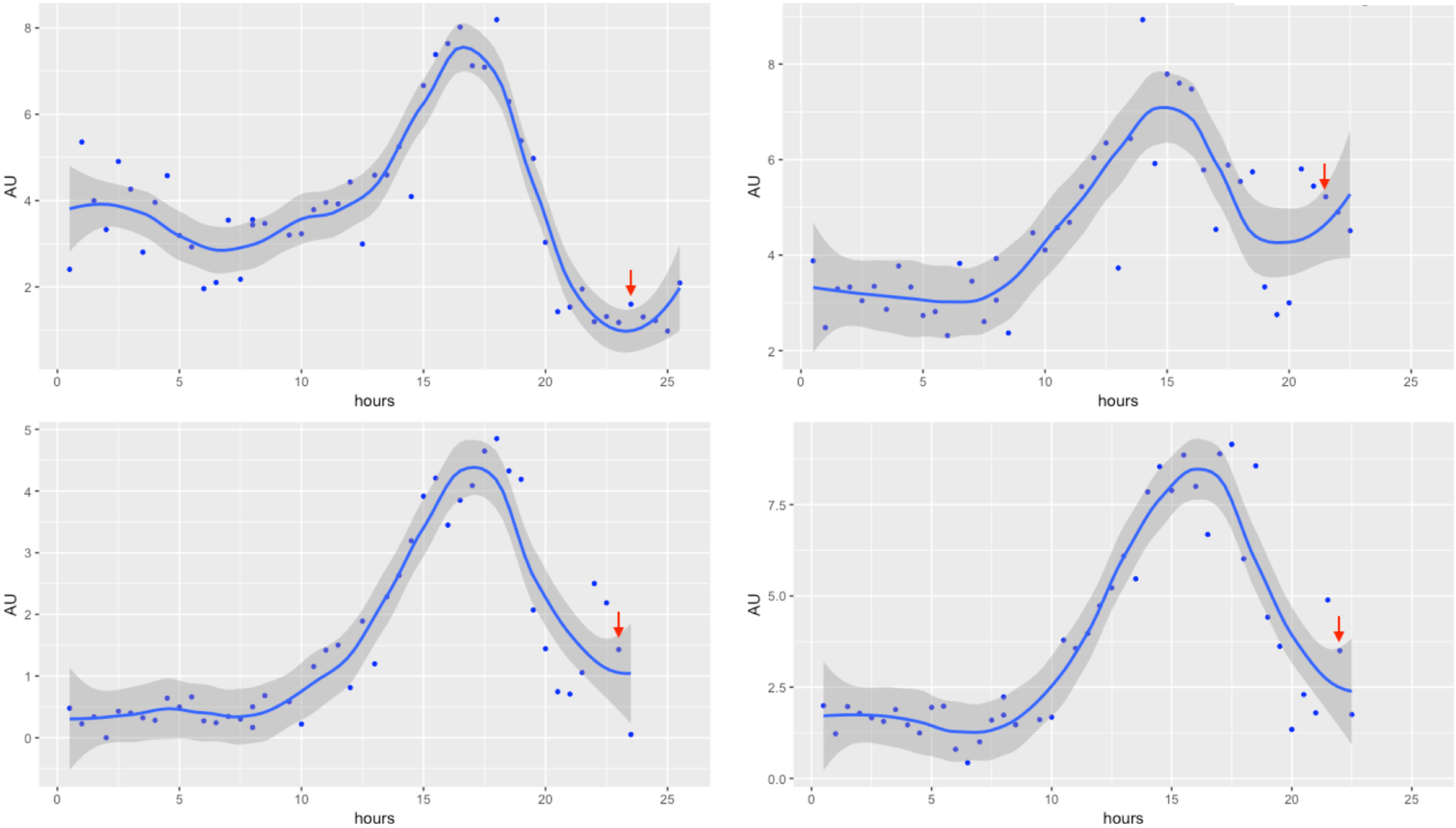
Time course of E2^V^F1 protein expression in REF52^E2vF1^ (clone 8) cells. Quiescent REF52^E2vF1^ (clone 8) cells, starved of serum for 48h, were released back into the cell cycle after addition of 10% serum. Cells were placed into the Vivaview incubator microscope and once the focus was fully stabilized (about 90 min) (t=0), images were recorded every 30 min in the DIC and YFP modes (20X objective) for 36h. E2^V^F1 fluorescent tracings for 4 representative cells are presented. AU: Arbitrary units of fluorescence. Red arrow: time of mitosis. Blue dots: values for each time point. Solid line: fitted curve (ggplot2: geom_point(), stat_smooth(method =“loess”, span = 0.4). Darker grey area: 95% confidence range for the fitted curve.

As the version of the protein reporter in clone 8 cells lacked the 3’UTR of the E2F1 gene and expressed a higher level of E2^V^F1 protein (Fig. 2A), it was possible that some aspect(s) of E2F1 protein regulation might not be properly represented with this reporter. In addition, there was the potential that ectopic expression of E2^V^F1 might alter the balance of complex formation between Rb and E2F1 in a cell and thereby affect our measurements (Hofmann et al. 1996; Campanero and Flemington 1997). To examine these possibilities, and also calibrate our protein reporter against an E2F activity reporter (E2F^act^) that we had recently characterized (Dong et al. 2018), we took advantage of a human mammary epithelial cell (h-Tert HME^E2Fact^) clone P1 that we had recently derived. We generated Rb- derivatives of this cell clone using CRISPR-Cas9 editing (HME^E2Fact^, Rb-), and selected a single cell clone for the rest of our experiments (clone 0). We then introduced the E2^V^F1-3’UTR reporter construct, driven by the human E2F1 promoter (Fig. 1B) into HME^E2Fact^ Rb+ (clone P1) or HME^E2Fact^ Rb- (clone 0) cells and isolated single cell clones from the respective transduced populations. To quantify fluorescent signals in live cultures, we selected one representative each from our Rb+ and Rb- clones, clone R15 (HME^E2Fact^, Rb+, E2^V^F1-3’UTR) and clone H4 (HME^E2Fact,^ Rb-, E2^V^F1-3’UTR) respectively. Both clones expressed comparable amounts of E2^V^F1 protein to that found in REF52^E2vF1-3’UTR^ clone D cells (Fig. 5, SupFig. 1). Live imaging of R15 and H4 cells, which had been starved of growth supplements for 48h and then switched to full medium was performed over a 40h period. We plotted the respective fluorescence intensities every 30 minutes (Fig. 6) as described earlier for REF52 clone 8 cells. In both HME H4 and R15 cells, the E2^V^F1 signal dynamics followed roughly the same kinetics as that observed in cells expressing the E2^V^F1 reporter lacking the 3’UTR (Fig. 4). However, in several cases, we noticed a slightly more complex dynamic behavior as revealed by a bi-phasic or non-monotonic increase of E2^V^F1 intensity before reaching its maximum amplitude. Furthermore, the signal returned to basal levels a bit sooner before cell division in cells expressing the reporter containing the 3’UTR, consistent with the idea that miRNAs specifically induced by c-myc and E2F1 bind the 3’UTR of the E2F1 gene to help sharpen the reduction of E2F1 protein before cell division (O’Donnell et al. 2005). Although there were no drastic differences in the kinetics of E2^V^F1 expression in HME H4 cells that lacked Rb, the overall intensity of the signal was reduced in these cells compared to single Rb+ R15 cells, despite the fact that unsynchronized and proliferating cells in the two clones express similar steady-state amount of E2^V^F1 protein as detected by immunoblotting (Fig. 5).

**Figure 5:**
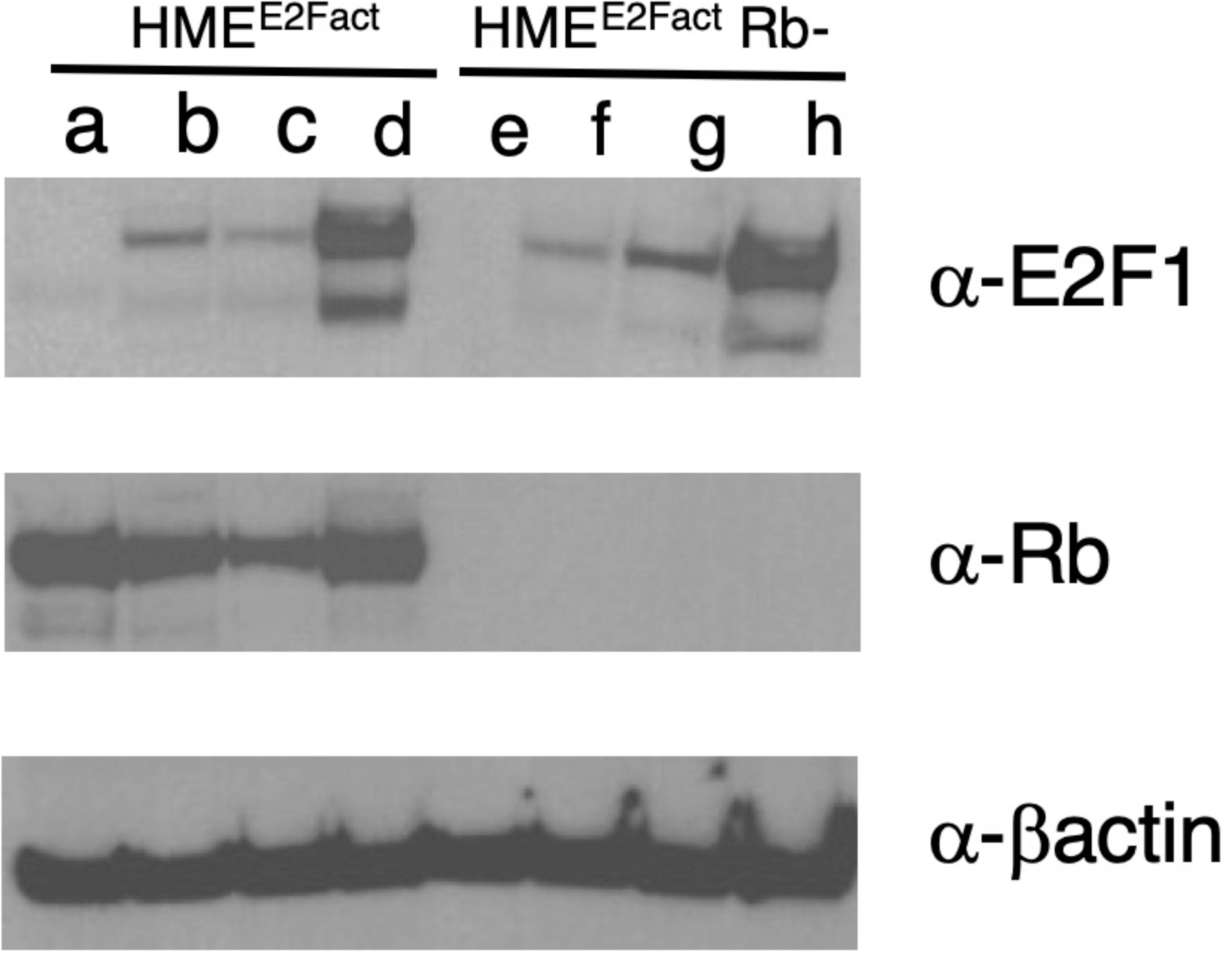
Rb+ and Rb-HME cells clones expressing both E2Factivity and protein reporters. Viral stocks of the E2^V^F1-3’UTR construct were used to infect a human mammary epithelial cell clone expressing the E2F activity reporter (HME^E2Fact^) (Dong et al. 2018) or a CRISPR-derived subclone harboring an Rb deletion (HME^E2Fact^ Rb-, clone 0). After selection, single cell clones were derived, and cell extracts were probed for expression of E2^V^F1, Rb and β actin respectively. (a) HME^E2Fact^ Rb+; (b) HME^E2Fact^ Rb+, E2^V^F1-3’UTR (polyclonal); (c) HME^E2Fact^ Rb+, E2^V^F1-3’UTR (clone R11); (d) HME^E2Fact^ Rb+, E2^V^F1-3’UTR (clone R15); (e) HME^E2Fact^ Rb- (clone 0); (f) HME^E2Fact^ Rb-, E2^V^F1-3’UTR (polyclonal); (g) HME^E2Fact^ Rb-, E2^V^F1-3’UTR (clone H1); (h) HME^E2Fact^ Rb-, E2^V^F1-3’UTR (clone H4).

**Figure 6:**
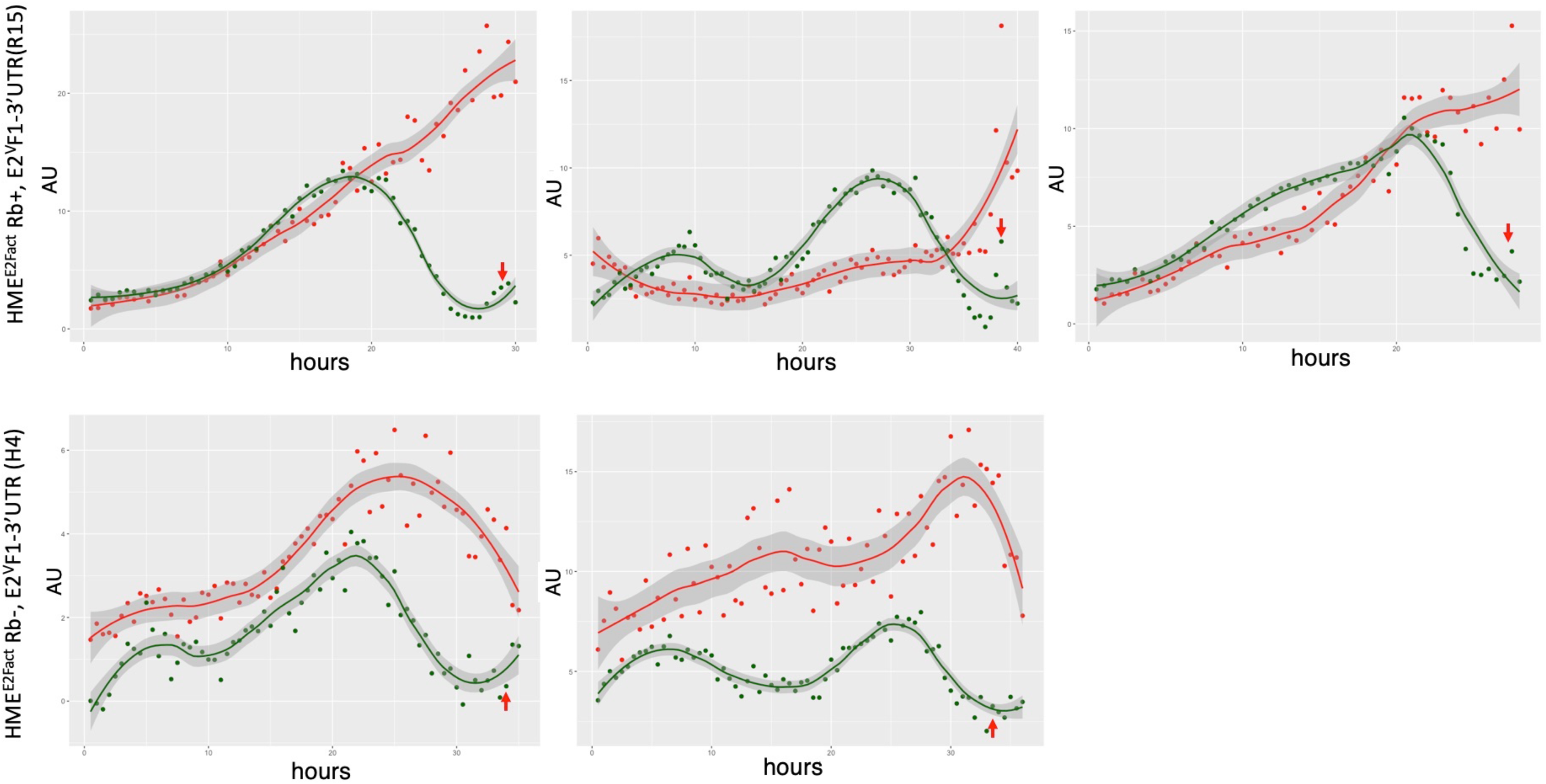
Time course of E2^V^F1 protein expression in HME cells. HME^E2Fact^ Rb+, E2^V^F1-3’UTR (clone R15)or HME^E2Fact^ Rb-, E2^V^F1-3’UTR (clone H4) cells (top 3 and bottom 2 panels respectively) were driven to quiescence by culturing them in minimum medium for 48h. Cells were then released back into the cell cycle after addition of full growth medium. Cells were imaged under DIC, RFP and YFP modes every 30 min for 40h with the Vivaview incubator microscope (20X objective) as described under Fig. 5. Representative time course tracings of E2F activity and E2^V^F1 protein reporters in singles cells. AU: Arbitrary units of fluorescence. Red arrow: time of mitosis. Solid lines: fitted curves for activity (red dots) and E2^V^F1 (green dots) reporters (ggplot2: geom_point(), stat_smooth(method =“loess”, span = 0.4). Darker grey area: 95% confidence range for fitted curves.

The overall kinetics of the E2F^act^ and E2^V^F1 reporters appear to track well during the cell cycle, although there are some differences. As we had noticed in our previous study (Dong et al. 2018), a strong signal from the E2F^act^ reporter can persist in cells at the time of cell division (Fig. 6, and data not shown), although its amplitude eventually decreases in daughter cells before it rises again. In contrast, the E2^V^F1 signal always returns to basal or low levels before cells undergo mitosis. Lastly, there is no strict correlation between the maximum amplitude values observed for the E2F^act^ and the E2^V^F1 reporters. This is likely due to the fact that the E2F^act^ reporter informs on the net activity of all E2F activators in the cell, whereas the E2^V^F1-3’UTR reporter is specific to E2F1 protein expression.

## Discussion

Over the last few years, we have assembled a versatile tool-set of reporters that inform on the kinetics of activation of E2F1 at the level of transcription, functional activity and now protein expression in live cells. These reporters can be followed in tandem (Venus vs. mCherry), which gives a unique opportunity to uncover more complex rules of regulation that cannot be captured by one reporter alone. So far, we have characterized and validated them in the context of cell proliferation, but they could be equally informative in studying the behavior and role of E2F1 in apoptosis, differentiation, and metabolism (Blanchet et al. 2011; Shats et al. 2017; Iaquinta and Lees 2007).

Our most recent E2F1 reporters, which capture the dynamic expression of E2F1 protein, may offer the most information as they reflect different levels of regulation. The reporter constructs are under the control of the mouse or human E2F1 proximal promoter (Dong et al. 2014), and also differ in including or excluding the E2F1 3’UTR. They encode a fluorescent fusion protein (E2^V^F1) (Fig. 2), consisting of Venus embedded within the coding region of E2F1, which is detected as a doublet, with the top band corresponding to the expected size for the fusion protein. To avoid disrupting the balance between functional E2F activators and repressors in cells that ectopically expressed E2^V^F1, the winged-helix DNA binding domain of E2F1 was deleted and replaced with Venus. The fact that this domain is equivalent to a region in E2F4 shown to make critical contacts with DNA (Zheng et al. 1999), and that the E2F1 R166H mutation, which resides in the corresponding region of E2F1 abrogate DNA binding by mutant E2F1 (Yu et al. 2011), strongly suggests that E2^V^F1 has no DNA binding activity. Although we have not tested DNA binding directly, ectopic expression of E2^V^F1 from the constructs containing the E2F1 3’-UTR has no discernable effect on normal cell behavior. We failed to see any changes in the rate of proliferation, viability or susceptibility to genotoxic treatment. However, we did note that REF52 ^E2vF1^ clone 8 cells, which express a high amount of the fusion were more sensitive to trypsinization and freezing conditions. These cells needed to be on a strict passaging schedule to avoid crisis in recovery or slow growth with signs of senescence after passaging. Whether this behavior is due to a slight dominant negative effect of the over-expression of E2VF1 has not been investigated.

Detection of the E2^V^F1 fluorescent signal is largely restricted to the nucleus, mirroring the localization of endogenous E2F1 protein (Magae et al. 1996). The biggest difference between the two reporter versions is that the reporter lacking the 3’UTR expresses a much higher amount of E2^V^F1 than the reporter containing the 3’UTR sequence. This differential expression was observed both in polyclonal populations and single cell clones of REF52 cells transduced with the reporters, indicating that the presence of the 3’UTR selectively reduces the overall transcription of the reporter construct, or more likely, the translation/accumulation of the fusion protein. It has been reported that the E2F1 3’UTR contains functional miRNA sites that bind miRs from the miR-20a and miR-17-92 cluster leading to a sharp decrease in E2F1 protein during the cell cycle as part of a delayed negative feedback loop (O’Donnell et al. 2005; Pickering, Stadler, and Kowalik 2009). Consistent with this explanation, reduction of E2^V^F1 signal to baseline level at the single cell level happens sooner and/or more often before the time of mitosis when the reporter contains the 3’UTR.

Time course trajectories of the E2^V^F1 fluorescent signal obtained from single cells freshly released into the cell cycle follow closely the pattern observed for endogenous E2F1 protein levels, and track well with the tracings obtained for the E2F1 transcriptional and activity reporters (Dong et al. 2014). Of note, we did observe slightly more complex shapes for the activation kinetics tracings of the E2^V^F1-3’UTR reporter compared to those from the construct lacking the 3’UTR. However, before more cells are analyzed, we cannot rule out that the observed differences arise from measurement fluctuations inherent to the lower fluorescent intensity of the E2^V^F1-3’UTR reporter and/or from intrinsic differences in activation between the different cell types in which the measurements were performed.

Given that the Rb binding site in E2F1 (Helin et al. 1992) is preserved in E2^V^F1, we wondered whether Rb might impact the behavior or level of E2^V^F1 signal during the cell cycle. At the population level and in unsynchronized cells, the amount of steady-state E2^V^F1 protein did not appear to be specifically sensitive to the presence or absence of Rb (data not shown), nor was there a consistent connection between E2^V^F1 and Rb protein levels in cells expressing both proteins (Fig. 5). Similarly, there were no dramatic changes when tracings of the E2^V^F1 signal from single cells containing or lacking Rb were compared (Fig. 6). The lack of obvious differences in the E2^V^F1 trajectories in Rb+ r Rb- clones was somewhat surprising given that the Rb- cells grew at higher density, which could have been the result of an accelerated cell cycle progression. However, we found no clear evidence for that, an observation consistent with the lack of overproliferation observed for Rb- cells populating mice chimeric for Rb status (Hanahan and Weinberg 2011). Interestingly, we noticed that cells in our Rb- polyclonal or single-cell clones were smaller and more motile than their Rb+ counterparts. In addition, Rb- cells were more sensitive to depletion of growth supplements normally present in the medium and adopted a stereotypical cobblestone morphology when arrested (SupFig. 4)

Many aspects of E2F1 function(s) and or stability are regulated by post-translational modifications (Poppy Roworth, Ghari, and La Thangue 2015). In particular, it well established that DNA damaging agents induce E2F1 phosphorylation, resulting in stabilization and accumulation of E2F1 (Lin, Lin, and Nevins 2001). E2^V^F1 was similarly stabilized in cells treated with cisplatin. Although the effect was fairly modest in clone 8 cells, which already express a high level of E2^V^F1, there was a dose-dependent stabilization of the reporter protein in REF52 clone D cells which express a more physiological level of E2^V^F1. Interestingly, stabilization of the top band of E2^V^F1 was more pronounced, as the lower band likely lacks serine 31, the residue targeted by ATM in response to DNA damage (Lin, Lin, and Nevins 2001).

## Material and Methods

### E2F1 protein reporters

*E2*^*V*^*F1 DNA:* The coding sequence of Venus flanked in frame by two short DNA sequences encoding flexible linkers fl1(*N*-GGSGGSGGSGGST-*C)* and fl2 (*N*-SGGGGSGGGGSGGGGSGS-*C)* was inserted within the coding region of human E2F1 in place of the sequence encoding amino acid 153 to 174 of E2F1. *pQCXIP-mE2F1p::E2*^*V*^*F1*: The E2^V^F1 DNA was fused downstream of the proximal promoter (−1,165 to +123) of mouse E2F1 gene and the resultant fragment was ligated between the BamH1 and Xba1 sites of the pQXCIP vector (Clontech). *pQCXIP/N-m (or h)E2F1p::pE2*^*V*^*F1-3’UTR* : the 3’UTR from human E2F1 cDNA was ligated downstream of the E2^V^F1 DNA cassette and the new fragment was either ligated downstream of the mouse or human E2F1 promoter (Dong et al. 2014). The new intermediates were inserted between the BamH1 and Xba1 sites of pQXCIP (or pQCXIN) (Fig. 1B). Retroviral stocks corresponding to the various constructs after transfection into ecotropic and amphotropic packaging cell lines (Plat-E or Plat-A cells respectively (Morita, Kojima, and Kitamura 2000)). The retroviral stocks were used to infect recipient REF52, WI-38 or hTert-HME^E2Fact^ (clone 1) cells (Dong et al. 2018). Puromycin (or neomycin when appropriate) was added to transduced cells for selection of a polyclonal population. Single cell clones were isolated by limiting dilution in the case of REF52 and hTert-HME^E2Fact^ polyclonal populations.

### Rb Gene editing

The human Rb gene locus was disrupted using the CRISPR-Cas9 system. Three optimized single-guide RNAs sequences (sgRNAs) were selected to target the region surrounding the start codon of human Rb. These sequences were subcloned into the lentiCRISPRv2 vector (Sanjana, Shalem, and Zhang 2014; Shalem et al. 2014) and lentiviral stocks were generated to independently infect human mammary epithelial HME^E2Fact^ (clone 1) cells (Dong et al. 2018). Bleomycin selection was applied and genomic DNA from polyclonal populations was isolated to screen for disruption of the Rb locus using the Surveyor Mutation Detector kit (IDT). Lack of Rb protein expression in cells targeted by Rb sgRNA (sequence 5’-GCGGTGCCGGGGGTTCCGCGG-3’) was further confirmed by immunoblotting, and a single cell clone (clone 0) was isolated from parental HME^E2Fact^ Rb- cells.

### Cell culture

REF52 cells (an immortal line of post-crisis Fischer rat embryo cells were grown in Minimum Essential Medium α (MEM α) (Gibco/Invitrogen) supplemented with 10% bovine growth medium (BGS, Hyclone/Thermo Scientific). For time-lapse microscopy and measurements of E2^V^F1 dynamics, ∼1 × 10^5^ REF52^E2vF1^ (clone 8) cells were seeded in p35 Mattek optic plates. Cells were synchronized to quiescence (G_0_) by culturing them in MEM α supplemented with 0.02% BGS (starvation medium) for 36 hours, after which the starvation medium was replaced with fresh medium containing 10% BGS and cells were moved to the Olympus VivaView incubator microscope. For cisplatin-induced DNA damage, ∼ 5 × 10^5^ REF52^E2vF1^ (clone 8) or REF52^E2vF1-3’UTR^ (clone D) cells were seeded in a 6 well tissue culture dish and grown in full medium overnight after which cisplatin (*cis*-Diaminedichloroplatimum III, Aldrich 479306) was added at varying concentrations for 24 hours and cell extracts were prepared. WI-38 cells (human fetal lung fibroblasts (Hayflick 1965)) (passage 18 through 32) were cultured in Eagle’s Minimum Essential Medium (EMEM) supplemented with 10% BGS. hTert-HME^E2Fact^ cells expressing the E2F^act^ reporter (Dong et al. 2018)) were routinely grown in full medium consisting of Mammary Epithelial Medium 171 (Medium 171; Thermo Fisher M171500) containing Mammary Epithelial Growth Supplement (MEGS; Thermo Fisher S0155). For time-lapse microscopy and measurements of E2^V^F1 dynamics, trypsinized cells were plated in p35 Mattek optic plates at 30% confluence and incubated overnight in full medium. To synchronize the cultures in G_0_, the medium was then replaced by 3 ml of Medium 171 lacking any supplement (starvation medium #1). After 24h, the starvation medium 1 was replaced by 3 ml of Medium 171 supplemented with 30 μ l of full medium (starvation medium #2). Cells were further incubated in this medium for 24h, at which point the starvation medium 2 was replaced by 3 ml of full medium and cultures were moved to the Olympus VivaView incubator microscope for live imaging.

### Live cell imaging

REF52 WT, REF52^E2vF1^ (clone 8), REF52 ^E2vF1-3’UTR^ (clone D) cells were imaged under brightfield or DIC illumination and YFP illumination (20X) to confirm expression and localization of the fusion protein. For time-lapse microscopy, quiescent cells growing in 35 mm p35 Mattek optic plates were released into the cell cycle and placed into the Olympus VivaView FL incubator microscope. Images were taken every 30 min for 36h using a 20×0.75 DIC Olympus UPlanSAPO 0.65 mm WD objective lens under DIC illumination and YFP 25% illumination (150 msec). Quiescent HME^E2Fact^ Rb+, E2^V^F1-3’UTR (clone R15) and HME^E2Fact^ Rb-, E2^V^F1-3’UTR (clone H4) cells were released into the cell cycle and placed into Olympus VivaView FL incubator microscope. Images were taken every 30 min for 40h using a 20×0.75 DIC Olympus UPlanSAPO 0.65 mm WD objective lens under DIC illumination (30 msec), RFP 25% illumination (300 msec), YFP 25% illumination (1000 msec) (binning 2). For time course fluorescence quantification, images were analyzed in ImageJ and values were determined as previously described (Dong et al. 2014).

### Fixed cell imaging

HME^E2Fact^ Rb-, E2^V^F1-3’UTR cells were plated on 12mm #1.5 coverslips pre-treated with a 0.1% gelatin solution and placed in single wells of a 6-well plate containing 3 ml of complete growth medium. After 24h, cells were fixed in 4% PFA (Image-IT™ ThermoFisher) according to the manufacturer’s recommendations and permeabilized in 0.3% Tween 20 for 5 min. The coverslips were then mounted on slides using Prolong Diamond with DAPI (ThermoFisher). Confocal images were obtained on the Zeiss 780 inverted confocal microscope (63X/1.4 NA Oil Plan-Apochromat DIC) in the 405 nm (DAPI), 514 nm (EYFP) and 561 nm (mcherry) channels respectively.

### Western blot analysis

Cells of interest were harvested, and protein extracts were obtained after cell lysis in RIPA buffer supplemented with a cocktail of protease inhibitors. Protein concentrations were determined using the Thermo Fisher BCA assay (Pierce). Equal protein amounts were run on Bolt 4-12% Bis-Tris Plus gels (Invitrogen) and transferred to polyvinylidene membranes by electroblotting. Membranes were blocked with 5% nonfat dried milk/TBST buffer (Blotto) for an hour, incubated for an hour with primary antibody, washed with TBS and 0.05% Tween (TBST), and then incubated again with secondary antibody in Blotto solution. Detection was performed via the LumiGLO Peroxidase Chemiluminescent Substrate Kit (Cell Signaling). The same membranes were re-probed after stripping to control for equal loading using β -actin as a control. Antibodies: *E2F1*: anti-hE2F1 3742S (1:1000, Cell Signaling); *Rb*: anti-Rb clone G3-245 (1:250, BD Biosciences); β *-actin*: anti-β -actin 8H10D10 (1:1000, Cell Signaling); Secondary antibodies: anti-Rabbit or anti-Mouse Ig-conjugated with HRP (1:1000, Cell Signaling).

## Acknowledgments

We thank Y. Gao for his help with time-lapse and confocal microscopy at the Duke Light Microscopy Core Facility. This research was supported by a grant from NIH (1R01-GM106107) (L.Y and B.M-P.), and funds from the School of Medicine at Duke University (B.M-P).

## Author contributions

B.M-P., P.D., G.Y. and L.Y. developed the concept of the paper. B.M.-P. designed the research approach and performed experiments with the help of P.D., B-T.P., C.I., C.H. B.M-P. analyzed the data; B.M-P., P.D, G.Y. and L.Y. interpreted the results and wrote the manuscript with contributions from C.I.

