## Supplemental Figures for "Quantifying E2F1 protein dynamics in single cells"

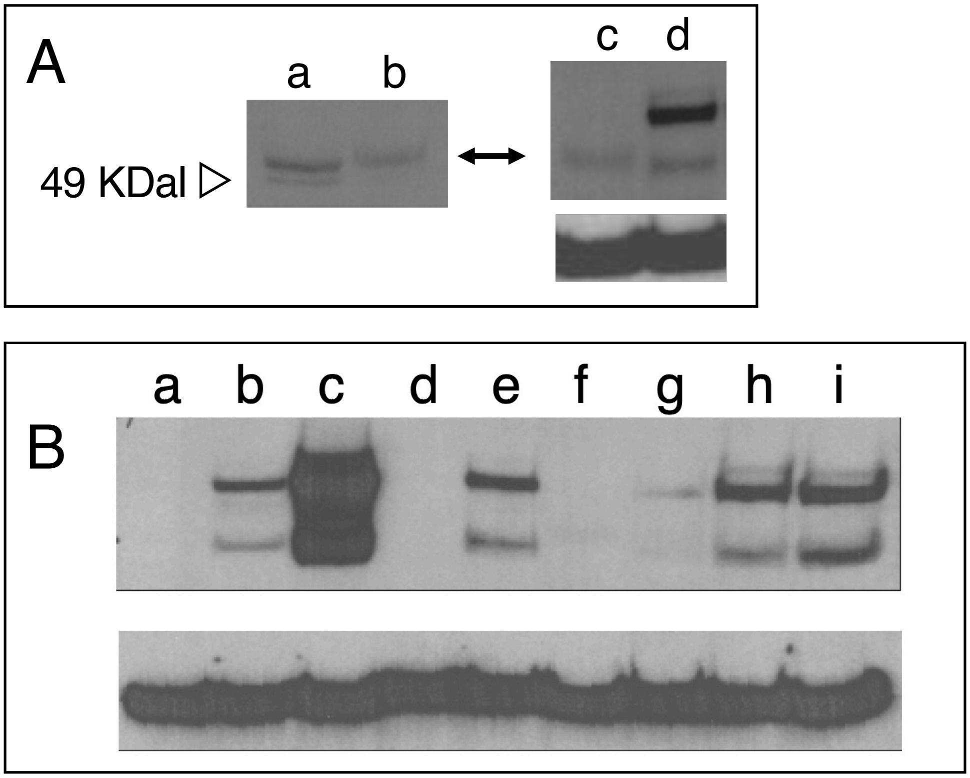


**Supplementary Figure 1: Detection of E2$^{V}$F1 in cell lines**

**A.** Cell extracts prepared from cells grown under normal conditions were probed for expression of E2$^{V}$F1. Top left and right panels: anti-E2F1 antibody. Left panel: Blot washed under less stringent conditions. Bottom panel on the right: anti β-actin antibody. (a) REF52 cells; (b) HME cells; (c) HME$^{E2Fact}$ (d) HME$^{E2Fact}$ Rb+, E2$^{V}$F1-3'UTR clone R1.

**B**. Cell extracts prepared from cells grown under normal conditions were probed for expression of E2$^{V}$F1. Top panel: anti-E2F1 antibody. Bottom panel: anti β-actin antibody. (a) REF52 cells; (b) REF52$^{E2vF1-3'UTR}$ clone D; (c) REF52$^{E2vF1}$ clone 8; (d) WI-38 cells; (e) WI-38$^{E2vF1-3'UTR}$ (polyclonal); (f) HME$^{E2Fact}$; (g) HME$^{E2Fact}$ Rb+, E2$^{V}$F1-3'UTR (clone R1); (h) HME$^{E2Fact}$ Rb-, E2$^{V}$F1-3'UTR (clone H1); (i) HME$^{E2Fact}$ Rb-, E2$^{V}$F1-3'UTR (clone H4).

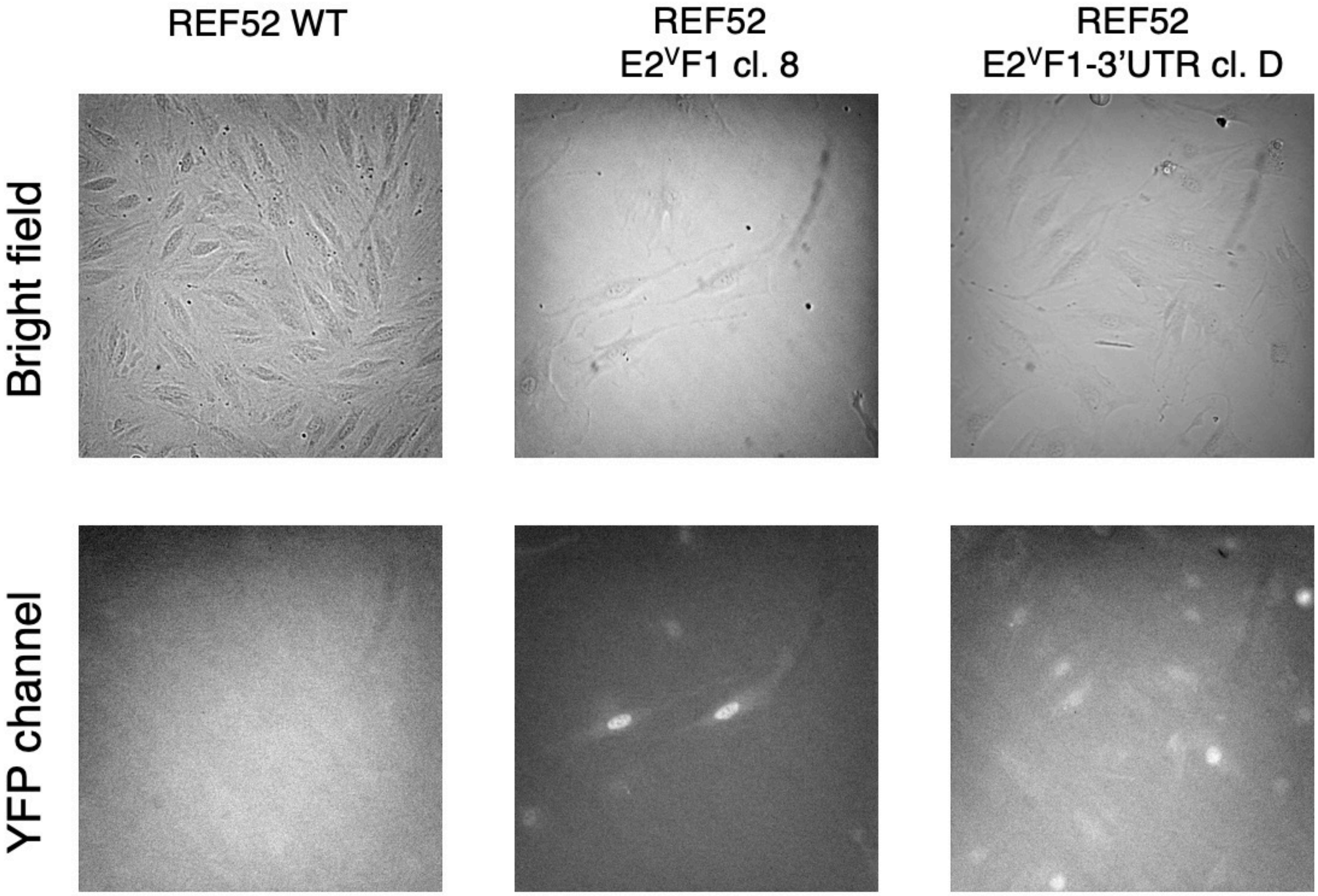


**Supplementary Figure 2: Live detection of E2$^{V}$F1 in single cells**.

Images taken under brightfield and YFP illumination at 20X of growing REF52 fibroblasts, REF52$^{E2vF1}$ cl. 8 and REF52$^{E2vF1-3'UTR}$ cl. D fibroblasts. The E2$^{V}$F1 fluorescent signal is largely localized to the nucleus of cl. 8 and cl. D cells, but the intensity is significantly weaker in cl. D cells.

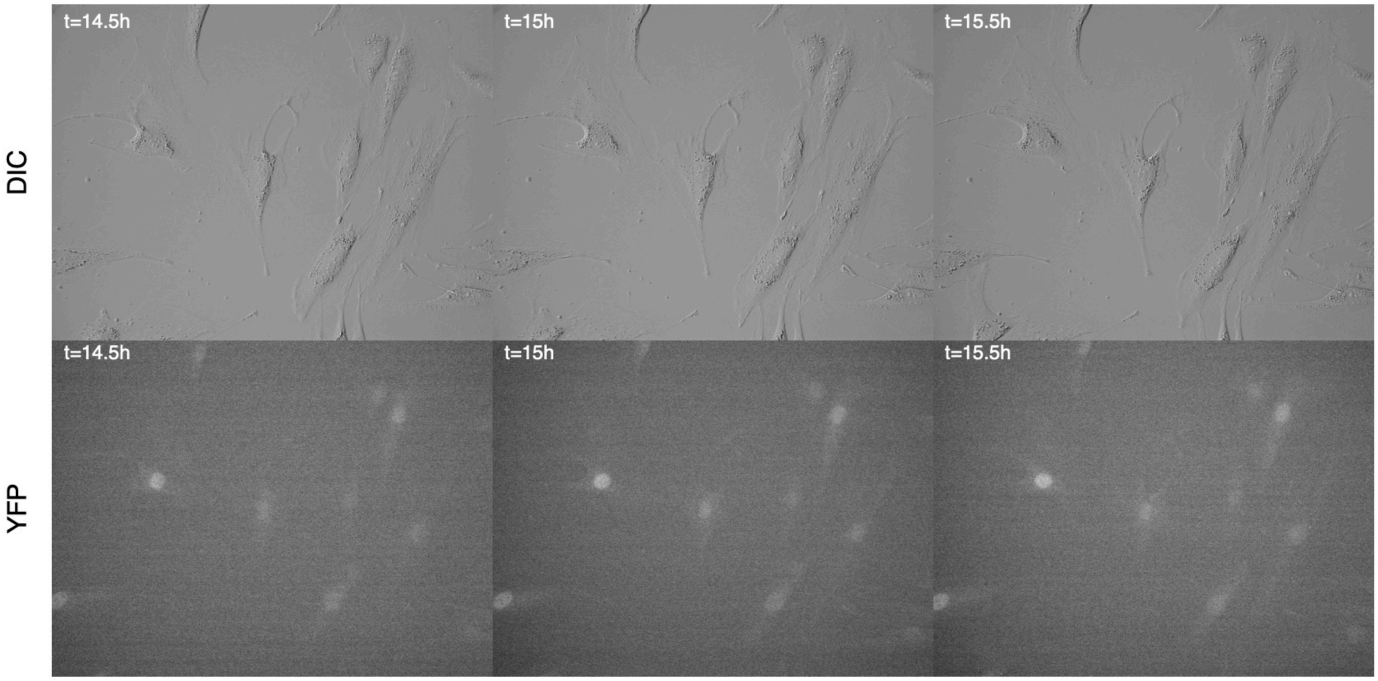


**Supplementary Figure 3: Dynamic expression of E2$^{V}$F1 expression**

Quiescent REF52$^{E2vF1}$ (clone 8) cells were starved of serum for 48h and released back into the cell cycle after addition of 10% serum). Live cell imaging was performed every 30 min in the Vivaview incubator microscope under DIC and YFP illuminations (20X objective) every 30 min. Selected images are shown for time t=14.5, 15 and 15.5h

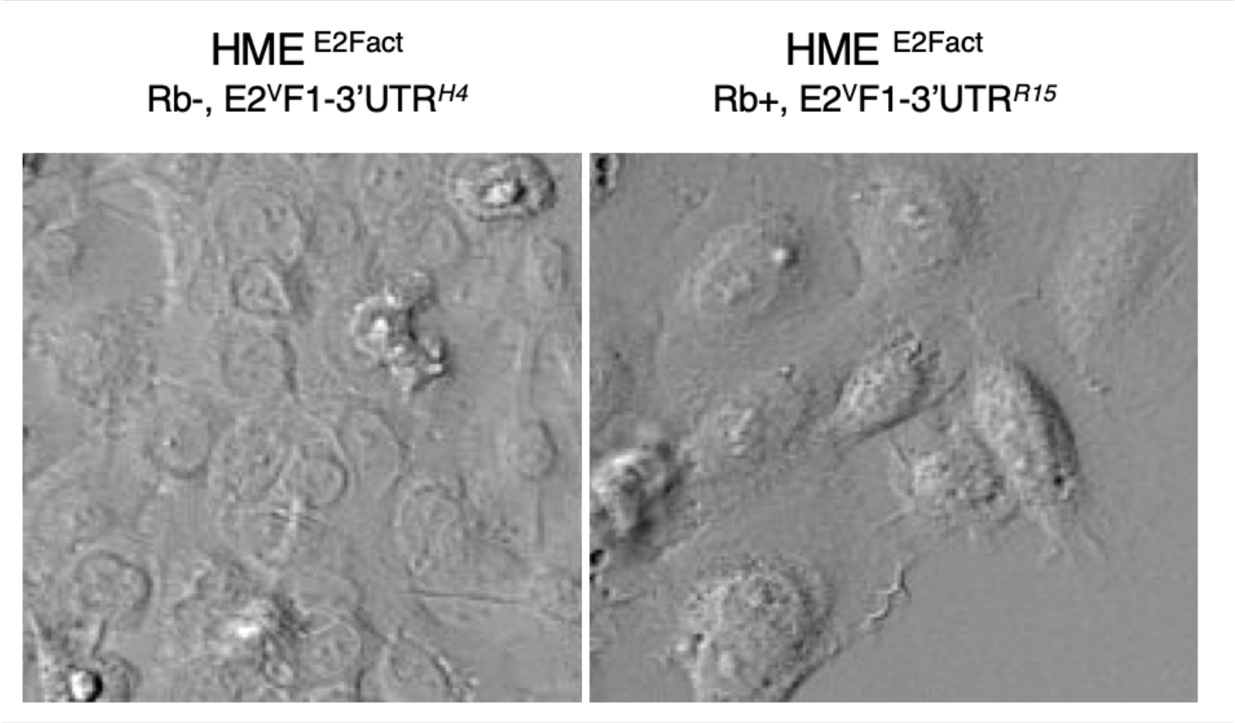


**Supplementary Figure 4: Morphology of HME clones arrested by growth signal deprivation.**

HME$^{\mathbf{E2Fact}}$ Rb-, E2$^{V}$F1-3’UTR cells (clone H4) (left panel) and HME$^{\mathbf{E2Fact}}$ Rb+, E2$^{V}$F1-3’UTR cells (clone R15) (right panel) were driven to quiescence by culturing them in minimum medium for 48h. Cells were then imaged under DIC, on the Vivaview incubator microscope (20X objective).
